# A functional interaction between TDP-43 and USP10 reveals USP10 dysfunction in TDP-43 proteinopathies

**DOI:** 10.1101/2024.01.23.576828

**Authors:** A Marrero-Gagliardi, J Noda, M Zanovello, G Gerenu, JM Brito Armas, A Bampton, P Torres, H Hernández-Eguiazu, S Moragón, F Pellegrini, C Pérez Hernández, F Fumagallo, L Taoro-González, R Muñoz de Bustillo Alfaro, AL Brown, G Quinet, P Andrés-Benito, I Ferrer, A Acebes, R Freire, VAJ Smits, FJ Gil-Bea, MJ Keuss, M Portero-Otin, T Lashley, P Fratta, A Acevedo-Arozena

## Abstract

Amyotrophic lateral sclerosis (ALS) and frontotemporal dementia (FTD) are fatal neurodegenerative disorders characterised by the progressive degeneration of specific neurons, that are defined by the appearance of TDP-43 pathology leading to TDP-43 cytoplasmic aggregation coupled with its nuclear loss. Although the causes of TDP-43 pathology in TDP-43 proteinopathies remain unclear, stress response may play a significant role, with some TDP-43 co-localizing with stress granules (SG). The ubiquitin-specific protease 10 (USP10) is a critical inhibitor of SG assembly. Here, we identify a new functional interaction between TDP-43 and USP10, with both proteins modulating different key aspects of the biology of the other. Adding to their functional connection, we assign a new function to USP10 as a modulator of alternative splicing, sharing a subset of splicing targets with TDP-43. Critically, we found that USP10 levels can increase in postmortem tissue from ALS and FTD patients and that USP10 can ameliorate TDP-43 mediated toxicity *in vivo* in an animal model, overall suggesting a new role for USP10 in TDP-43 proteinopathies.

## INTRODUCTION

Amyotrophic lateral sclerosis (ALS) is a fatal neurodegenerative disease characterised by the progressive degeneration of motor neurons in the cerebral cortex, brainstem and spinal cord, with a rapid progression from diagnosis to death of typically less than five years (1). ALS shares genetic and pathological features with the neurodegenerative disorder frontotemporal dementia (FTD). Approximately 20% of people with ALS meet the clinical criteria for a diagnosis associated with FTD (2). Both diseases share genetic, clinical and pathological features, with each representing the extremes of a broad disease spectrum (1,3). TDP-43 is a ubiquitously expressed and predominantly nuclear RNA/DNA binding protein encoded by the *TARDBP* gene that is involved in multiple aspects of RNA processing through its RNA binding capacity (4,5). TDP-43 plays a central role in ALS and FTD pathogenesis as a major component of the proteinaceous inclusions that pathologically define the great majority (>95%) of ALS cases and that are present in up to 50% of FTD cases (1,6). TDP-43 pathology occurs when TDP-43 is depleted from the nucleus and is sequestered as hyperphosphorylated insoluble aggregates in the cytoplasm, nucleus and/or dystrophic neurites of affected neurons. These features are also present in all cases of the dementia entity limbic-predominant age-related TDP-43 encephalopathy (LATE) and up to 50% of Alzheimer’s disease (AD) cases, which along with other disorders exhibiting similar TDP-43 pathology are collectively known as TDP-43 proteinopathies (7). The cytoplasmic aggregation of TDP-43 confers a loss of nuclear function, causing pleiotropic defects including splicing dysregulation such as the expression of toxic cryptic exons (8). Additionally, a possible toxic gain-of-function due to cytoplasmic mislocalization can contribute to the further nuclear clearance of TDP-43 and other RNA binding proteins (RBPs) (9–12).

TDP-43 and other RBPs involved in ALS, such as FUS, are components of stress granules (SG) - membraneless macromolecular ribonucleoprotein cytoplasmic condensates that are formed in response to different cellular stresses. Although the function of SG is not fully understood, they likely promote cell survival under stress conditions by storing stalled translation components, including mRNAs, RBPs, ribosomal components and initiation factors (13–15). For their formation, SG are thought to require an increase in cytoplasmic free RNA concentrations to promote the condensation of the core SG protein G3BP1 (16,17). Interestingly, TDP-43 controls *G3BP1* levels through direct mRNA binding, modulating SG formation (18–21). A key protein in the regulation of SG assembly is the ubiquitin-specific peptidase 10 (USP10), which inhibits SG formation by negatively influencing G3BP1 cooperativity via direct binding, in an interaction with G3BP1 that is mutually exclusive with that of Caprin1, which promotes SG formation (16, 17, 22–24). USP10 is a multifunctional protein with several roles that could be dependent on its enzymatic deubiquitinase activity, including controlling the levels of critical proteins like p53 or AMPK (25) and ribosome quality control (26,27), or independent of this activity, such as the inhibition of SG formation (16,17).

Recently, USP10 has been associated with different neurodegenerative conditions, colocalizing with tau aggregates in AD patients and is thought to promote tau aggregation in neuronal cells exposed to stress (28). In a cellular model of TDP-43 proteinopathy, USP10 promotes the elimination of TDP43-positive SG upon proteasome inhibition, whilst USP10 downregulation promotes the accumulation of TDP-43 insoluble fragments in the cytoplasm (29). Furthermore, USP10 was identified as a modifier of C9Orf72 toxicity in *Drosophila* (30).

Here, using different cellular models of TDP-43 cytoplasmic mislocalization to search for aggregation suppressors, we found that USP10 inhibits the formation of TDP-43 cytoplasmic inclusions through a mechanism that requires USP10 interaction with G3BP1. In different cellular models, we went on to study the effects of USP10 in modulating other aspects of TDP-43 biology, including its expression and splicing functionality. Furthermore, we discovered that USP10 can, on its own, modulate transcriptome-wide alternative splicing, with a clear overlap with some TDP-43 splicing targets. Critically, we show that USP10 can rescue TDP-43 associated toxicity *in vivo* in a *Drosophila* model. Finally, we found that, in turn, TDP-43 can also control different aspects of USP10 biology, including its expression, which we determined is dysregulated in ALS and FTD patients, further implicating USP10 in the pathogenesis of TDP-43 proteinopathies.

## RESULTS

### USP10 controls TDP-43 aggregation

To test the effects of USP10 on TDP-43 cytoplasmic aggregation, the cardinal feature of TDP-43 proteinopathies, we first used a cellular model in HeLa cells transfected with a construct carrying a GFP-tagged TDP-43 harbouring mutations in its nuclear localization signal (NLS) (GFP_TDP-43^ΔNLS^), leading to its mislocalization and the formation of cytoplasmic aggregates (31). Co-transfecting GFP_TDP-43^ΔNLS^ with USP10 resulted in a major decrease in both the size and the percentage of cells with TDP-43 cytoplasmic inclusions when compared to the effects of GFP_TDP-43^ΔNLS^ alone (GFP-ΔNLS, Fig. 1A). USP10 overexpression levels were controlled via western blots (Fig. S1A). To ensure that these results were not dependent on cell type, we performed the same experiments in U2OS and SH-SY5Y cell lines, largely confirming the results from Hela (Fig. S1B, C). We also tested the effects of USP10 downregulation, co-transfecting the GFP_TDP-43^ΔNLS^ construct with siRNAs against *USP10*. Reassuringly, USP10 downregulation led to the opposite results obtained when USP10 was upregulated, causing an increase in the size of TDP-43 inclusions when co-transfected with GFP_TDP-43^ΔNLS^ (Fig. S1D).

**Figure 1.**
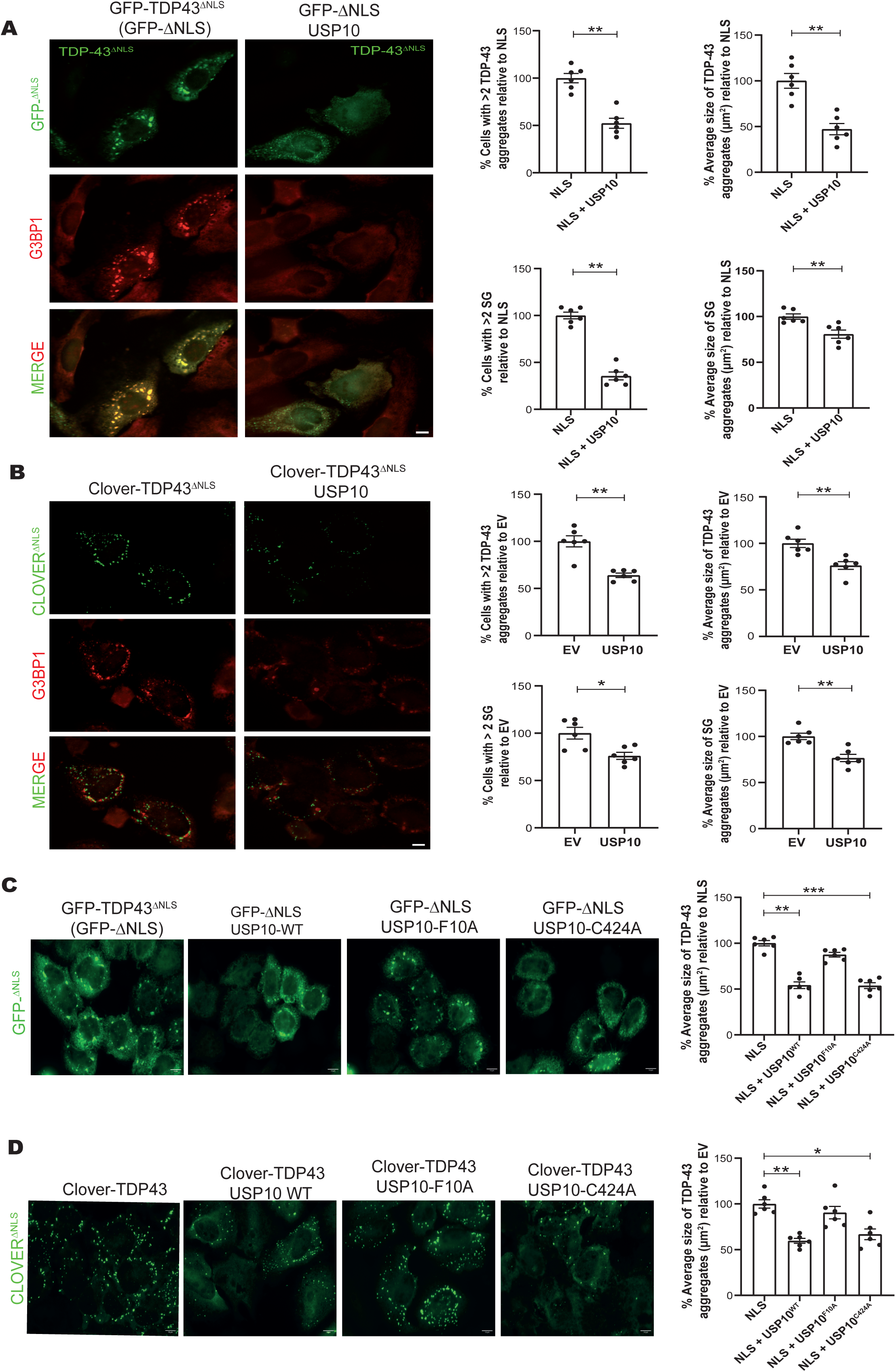
TDP-43 cytoplasmic aggregation is modulated by USP10 through its interaction with G3BP1. **(A)** Representative fluorescence images showing GFP_TDP-43 aggregates in HeLa cells 24 hours after co-transfection with GFP_TDP-43^ΔNLS^ and USP10 or an EV, co-labelled with anti-G3BP1 primary antibody in red marking SG. Scale bar: 10 µm. Quantification of the percentage of cells with more than two GFP_TDP-43^ΔNLS^ aggregates bigger than 1 µm; the average size of the TDP-43 aggregates; the percentage of cells with more than two SG bigger than 0.5 µm; and the average size of SG, comparing in all panels GFP_TDP-43^ΔNLS^ cotransfection with an EV (NLS group) with the co-transfection of GFP_TDP-43^ΔNLS^ and USP10 (NLS + USP10). For all figure panels, each datapoint corresponds to one independent experiment on which at least 50 cells per group were analysed. *p<0.05, **p<0.01, ***p<0.001 analysed by Mann-Whitney U test, with error bars showing the SEM for all panels. **(B)** Representative fluorescence images showing TDP-43 aggregates in green and SG labelled with G3BP1 in red, in U2OS cells 24 hours after endogenous Clover_TDP-43^ΔNLS^ induction with doxycycline and transfection with an EV or USP10 followed by induction of TDP-43 aggregation with 250 µM NaAsO_2_ for 80 minutes. Scale bar: 10 µm. Quantification of the average size and percentage of cells with more than two Clover_TDP-43 aggregates bigger than 0.5 µm after transfection with USP10 or EV, together with the size and percentage of cells with SG bigger than 0.5 µm. **(C)** Representative images of TDP-43 aggregates (green) in HeLa cells 24 hours after co-transfection with GFP_TDP-43^ΔNLS^ and either an EV, USP10^WT^, USP10^F10A^ or USP10^C424A^. Scale bar: 10 µm. **(D)** Representative images of TDP-43 aggregates (green) in doxycycline inducible TDP-43^ΔNLS^ U2OS cells 24 hours after doxycycline and 80 minutes after NaAsO_2_ (250 µM) treatment. Cells were transfected with either an EV, USP10^WT^, USP10^F10A^ or USP10^C424A^. Scale bar: 10 µm.

Given that TDP-43 and other RBPs associated with ALS can accumulate in SG (32,33) and knowing that USP10 inhibits SG assembly (16,17), we wondered whether the effects of USP10 on TDP-43 aggregation might be mediated by USP10’s role in regulating SG formation. Thus, in the same Hela cellular model transfected with GFP_TDP-43^ΔNLS^, we also quantified SG by immunocytochemistry using antibodies against G3BP1. As expected for a known inhibitor of SG formation, USP10 co-transfection with GFP_TDP-43^ΔNLS^ led to a clear reduction in SG size and a decrease in the number of cells with SG when compared with the co-transfection with the empty vector (EV) (Fig. 1A). In this model, the colocalization between TDP-43 cytoplasmic foci and SG was very high, as the expression of the cytoplasmic GFP_TDP-43^ΔNLS^ construct itself leads to SG formation (Fig. S1E).

Because of the almost complete colocalization between TDP-43 and G3BP1 in this experimental setting, it was not possible to distinguish between USP10’s roles as a generic SG formation inhibitor from its possible effects on TDP-43 aggregation. To directly address if USP10 could affect TDP-43 aggregation independently of its co-localization in SG, we turned to a different cellular model in which the endogenous *TARDBP* gene is edited to harbour a disrupted NLS and a fluorescent tag (TDP-43^ΔNLS^-Clover) expressed under a doxycycline-inducible system that lead to TDP-43 cytoplasmic aggregation under oxidative stress conditions (12). Under these conditions, SG and TDP-43 foci were both formed and were largely independent, allowing to assess the effects of USP10 specifically on endogenous TDP-43 aggregates that do not colocalize with SG (Fig. 1B and S1E). In this model, USP10 overexpression, when compared to the EV control, also led to a decrease in the percentage of cells with TDP-43 inclusions as well as in the number of TDP-43 inclusions per cell (Fig. 1B). USP10 overexpression levels were controlled via western blots (Fig. S1A). Similar results were also obtained in a third cellular model, on which SH-SY5Y cells were treated with oxidative and osmotic stress, again assessing endogenous TDP-43 inclusions (Fig S1F).

Thus, USP10 can modulate TDP-43 aggregation in all tested conditions, irrespective of the cause of aggregation and independently of its colocalization with stress granules.

### The effect of USP10 on TDP-43 cytoplasmic aggregation is independent of its deubiquitinase activity, but dependent on its binding to G3BP1

USP10’s deubiquitinase activity is critical for its involvement in regulating different cellular processes (24, 34–36), but not for the regulation of SG formation (16). Thus, to begin to determine the mechanisms by which USP10 inhibits TDP-43 aggregation, we used the same two distinct cellular models (the overexpression of GFP_TDP-43^ΔNLS^ in HeLa cells, as well as the inducible TDP-43^ΔNLS^-Clover U2OS model), to test the effects of a mutated version of USP10 devoid of its catalytic activity (USP10^C424A^) (37, 38). The expression of enzymatically inactive USP10^C424A^ led to a decrease in TDP-43 aggregation that was indistinguishable from that of USP10^WT^ in both cellular models (Fig. 1C, D), showing that USP10 effects on TDP-43 aggregation are not dependent of its deubiquitinase activity.

We then tested for a possible role of the interaction between USP10 and G3BP1 on USP10’s capacity to clear TDP-43 aggregates by using a USP10^F10A^ mutant that loses its ability to bind G3BP1 (24). As already published in other contexts (16), USP10^F10A^ overexpression, did not inhibited SG formation in our co-transfection model with GFP_TDP-43^ΔNLS^ (Fig. S1G). We also confirmed by immunoblot that all USP10 constructs were expressed at similar levels (Fig. S1H). Crucially, unlike USP10^WT^ or USP10^C424A^, USP10^F10A^ had no significant effects on TDP-43 aggregation in any of the models, suggesting that the inhibitory role of USP10 on TDP-43 aggregation is mediated by its interaction with G3BP1 (Fig. 1C, D). Overall, these results show that USP10 binding to G3BP1 is required for its inhibition of TDP-43 aggregation in a manner that is not dependent on TDP-43 colocalization with SG.

### USP10 enhances TDP-43 functionality on splicing

Next, we tested whether USP10 could affect TDP-43 functionality at the mRNA splicing level. First, we employed a widely used *CFTR* reporter minigene in which TDP-43 function is known to modulate the inclusion/exclusion ratio of *CFTR* exon 9 (39). We co-transfected the *CFTR* minigene together with USP10 in HeLa cells and assessed the percentage of exclusion (PSE) of *CFTR* exon 9. Our results show that USP10 overexpression enhance endogenous TDP-43 splicing function when compared to the EV (Fig. 2A).

**Figure 2.**
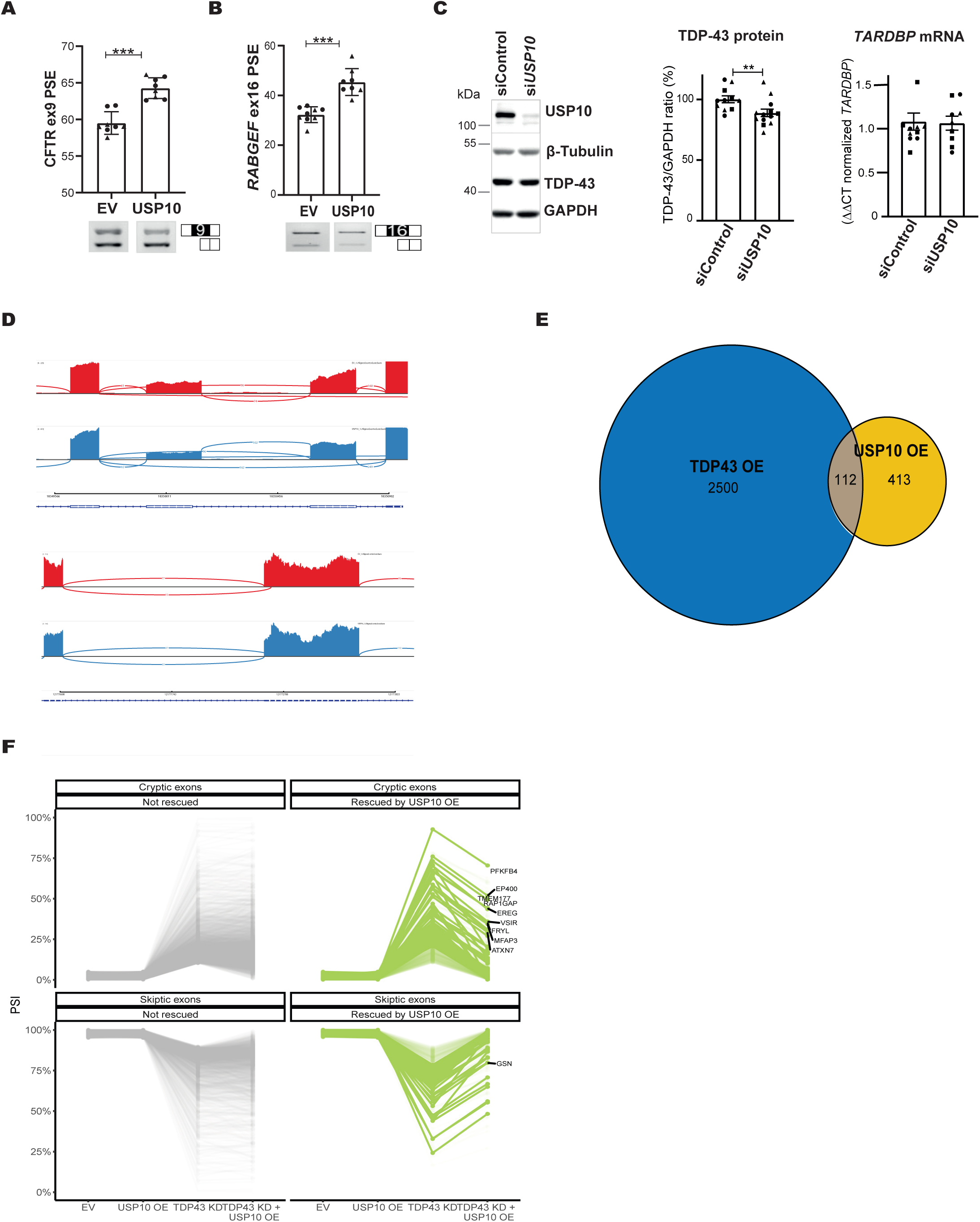
USP10 regulates alternative splicing and enhance TDP-43 splicing function. **(A)** Representative image and quantification of the PSE (percentage of splicing exclusion) of the *CFTR* exon 9 minigene. HeLa cells were cotransfected with the *CFTR* minigene and either EV or USP10. Following RT-PCR, the PSE was quantified by densitometry, comparing to the EV via one-way ANOVA. For all figure panels, each data point represents a different cellular well, with the different shapes representing independent experiments. *p<0.05, **p<0.01, ***p<0.001 and error bars represent SEM for all figure panels. **(B)** Representative image and quantification of the PSE of *RABGEF* exon 16. HeLa cells were transfected with either an EV or USP10. Following RT-PCR, the PSE was quantified and compared via one-way ANOVA. **(C)** Representative image of immunoblots in U2OS cells treated with siRNA against *USP10* (siUSP10) or siRNA against luciferase (siControl). TDP-43 levels were quantified via densitometry, using GAPDH as loading control, displaying the ratio of the siControl group as 100%. *p<0.05, by Mann-Whitney U-test. N=13, from 3 independent experiments. And quantification of the relative ΔΔCt expression levels of *TARDBP* transcript with *HPRT* control does not show a significant difference after USP10 downregulation when compared to siControl by Mann-Whitney U-test (p=0.3). N=10, from 3 independent experiments. (**D**) Sahimi plot examples of splicing events modified by USP10 overexpression. Top: *PKD1P4* skipping of exon 6. Bottom: *DAB2IP* splice site inside exon 11. **(E)** Venn diagram showing the overlap between splicing events shared between USP10 and TDP-43 overexpression. **(F)** Representation of the changes in the PSI of skiptic exons (bottom) and cryptic exons (top) that are rescued (green) or not (grey) by the overexpression of USP10 after TDP-43 downregulation in HeLa cells. A subset of the most changed splicing events is also annotated with the gene name.

In addition to the *CFTR* reporter minigene, we also assessed the PSE of endogenous exons known to be controlled by TDP-43 function, such as *RABGEF* exon 16 and *RWDD1* exon 2. In agreement with the minigene data, USP10 overexpression caused an increase in TDP-43 splicing function on both targets when compared to the EV controls (Fig. 2B and Fig. S2A). We obtained similar results in U2OS (Fig. S2B), demonstrating that, independently of the cell line, USP10 can positively modulate TDP-43 splicing function. This enhancement of TDP-43 splicing function upon USP10 overexpression occurs without affecting endogenous TDP-43 expression levels (Fig. S2C). However, acute USP10 downregulation in U2OS cells caused a small but significant reduction in endogenous TDP-43 protein when compared to the siRNA control (Fig. 2C), without affecting *TARDBP* mRNA levels. Thus, USP10 can modulate TDP-43 splicing functionality and protein expression levels.

### USP10 affects alternative splicing overlapping with events controlled by TDP-43

As we show that USP10 can enhance TDP-43 splicing function, we wondered whether USP10 could, on its own, affect transcriptome-wide alternative splicing. Thus, we performed RNA-sequencing (RNAseq) analysis on HeLa cells transfected with USP10 and compared to an EV control. Indeed, USP10 significantly affected the inclusion of around 500 alterative splicing events (Fig. 2D, E), with around half of those increasing the inclusion of the identified splicing events modified by USP10. Next, we wanted to compare the splicing events modified by USP10 with those modified by TDP-43. As we already showed for a couple of targets that USP10 can positively affect the splicing function of TDP-43, we compared the overall splicing changes modified by USP10 overexpression with those affected by TDP-43 overexpression in the same cellular setting. Remarkably, the comparison of the total of the 525 events regulated by USP10 with those modified by TDP-43 overexpression (2500), revealed a clear overlap between the splicing junctions modified by USP10 with a subset of those controlled by TDP-43, with around a quarter of the USP10 controlling splicing events being also modified by TDP-43 (Fig. 2E). To assess the directionality of the changes, we defined the events modified in opposite directions by TDP-43 in our cellular model; for this, we transfected TDP-43 in HeLa cells and compared the RNA-seq obtained from TDP-43 overexpression with downregulation of TDP-43 using an inducible shRNA model in HeLa cells (40). Combining both datasets, we identified around 1650 splice junctions changing in opposite directions between TDP-43 downregulation and overexpression, that we defined as *bona fide* TDP-43 targets in our cellular setting. Focussing on these 1650 targets, we show that around a quarter of these *bona fide* TDP-43 targets move in our USP10 overexpression setting in the same direction as TDP-43 overexpression (Fig. S2D), further supporting a role for USP10 in enhancing TDP-43 splicing function at the transcriptome-wide level. Altogether, these results show for the first time that USP10 can modify alternative splicing, with a clear correlation between some of the splicing events controlled by USP10 and a subset of those controlled by TDP-43.

To assess if USP10 could also enhance TDP-43 splicing function even under TDP-43 loss of function conditions, we turned to our inducible shRNA model in HeLa cells, quantifying the effects of USP10 overexpression under conditions of TDP-43 loss of function on transcriptome-wide alternative splicing via RNA-seq analysis. We separated the events that increased percentage of exon inclusion (PSI) upon TDP-43 silencing, which included cryptic exons, and those that decreased PSI after TDP-43 downregulation, including skiptic exons (41). Remarkably, USP10 overexpression was able to rescue a proportion of the TDP-43 dependent cryptic, skiptic and cassette exons that changed upon TDP-43 downregulation (Fig. 2F). Importantly, all these changes occurred without significantly affecting *TARDBP* silencing (Fig. S2E). Thus, USP10 can modify the expression of at least a subset of TDP-43 dependent cryptic and skiptic exons, counteracting some of the toxic effects of TDP-43 loss of function.

### USP10 is affected in TDP-43 proteinopathies

As TDP-43 proteinopathies are defined at the pathological level by cytoplasmic TDP-43 inclusions, we proceeded to assess USP10 in patient post-mortem material. At the neuropathological level, we performed immunohistochemistry with antibodies against USP10 and phosphorylated TDP-43 (pTDP-43) in paraffin-fixed post-mortem samples from the frontal cortex of FTD patients, separating them by the different types of TDP-43 pathology (FTLD-TDP type A, B or C), which are classified based on the morphology and location of TDP-43 inclusions in the affected cortical areas, and whether they occur in conjunction with ALS (42,43). In agreement with a previous finding from spinal cord of ALS patients (29), we did not find any co-localization between USP10 and pTDP-43 aggregates by immunofluorescence analysis, but we nevertheless verified the presence of the characteristic pTDP-43 staining in all FTLD-TDP pathology subtypes (Fig. 3A). Interestingly, in the frontal cortex of FTLD-TDP type B patients, and to a lesser extent also in FTLD-TDP type A samples, we found increased cytoplasmic immunoreactivity of USP10 in some neurons showing characteristic USP10 staining (Fig. 3B).

**Figure 3.**
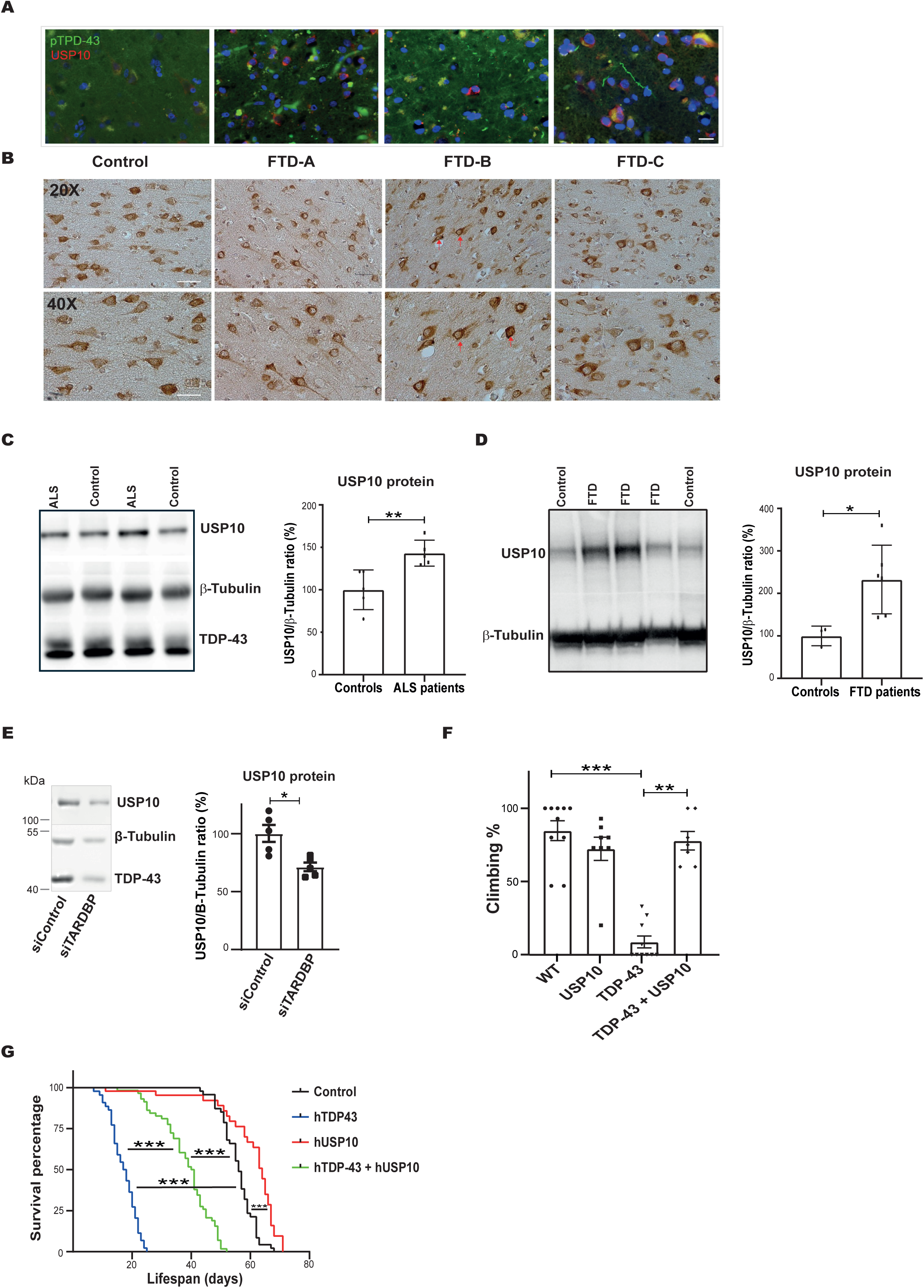
USP10 is affected in TDP-43 proteinopathies and can ameliorate TDP-43 toxicity *in vivo.* **(A)** Representative immunofluorescence and **(B)** immunohistochemistry images of post-mortem slices of frontal cortex samples from FTD patients. In (A), primary anti-pTDP-43 labelled in green, anti-USP10 in red, and nuclei in blue (DAPI), showing the merged images from the three channels. In (B) primary anti-USP10 revealed with DAB. Red arrows show increased USP10 immunoreactivity in the cytoplasm of some neurons. Scale bars: 50 μm. **(C)** Representative image of immunoblot and quantification of USP10 in protein lysates of motor cortex from 5 ALS patients and 5 controls and **(D)** frontal cortex from FTD 6 patients and 3 controls. For both panels, USP10 bands were quantified by densitometry, using β-Tubulin as loading control, normalizing to the average of each of the control groups (100%), displaying the percentage of the USP10/β-Tubulin ratio and comparing by Mann-Whitney U-test. (**E**) Representative image of immunoblot and quantification of SH-SY5Y downregulated for 72 h with either siRNA against *TARDBP* or a negative control (siControl). Quantification of USP10 bands by densitometry was normalized to their loading control (β-Tubulin). The quantification shown corresponds to a N=3, from one experiment, corroborated by another independent experiment, compared by T test to control. **(F)** *Drosophila* locomotor activity for the selected genotypes tested on day 7, quantified by the ability to climb an 8 cm mark inside a tube, normalised by the performance of the control group (100%), compared by Kruskal Wallis test. **(G)** Percentage of survival of the selected *Drosophila* genotypes of 50 adult female flies for each strain compared by Mantel-Cox. *< 0.05; **p<0,01 and ***p<0,001, with error bars show SEM for all figure panels.

Next, we assessed USP10 protein levels by immunoblotting in lysates from ALS and FTD patients cortex, finding a significant increase in USP10 levels in both sets of patients versus controls (Fig. 3C, D). Separating the FTD patients by the type TDP-43 pathology, uncovered a particular increase in USP10 levels on those with FTLD type B TDP-43 pathology, the subtype most commonly associated with ALS, characterised by the presence of TDP-43 positive neuronal cytoplasmic inclusions in all cortical layers (Fig. S3A). Thus, USP10 levels and subcellular localization are altered in the cortex of different TDP-43 proteinopathies.

As USP10 levels increased in TDP-43 proteinopathies patients, we sought to assess the levels of USP10 in different cellular models. We modulated TDP-43 levels by either overexpression or downregulation in different cell lines and assessed possible effects on endogenous USP10 levels at the protein and transcript levels. TDP-43 overexpression did not significantly affect endogenous USP10 levels in HeLa or other cell lines (Fig. S3B). However, acute downregulation of TDP-43 levels by siRNA in HeLa or SH-SY5Y cells led to a significant reduction in endogenous USP10 protein (Fig. 3E and S3C), that was also reflected at the transcript level and was confirmed in the RNA-seq data from HeLa cells with an inducible downregulation of *TARDBP* via shRNA (Fig. S3D). On the other hand, chronic downregulation of *TARDBP* in an inducible human induced pluripotent stem cell i3 model differentiated onto motor neurons (44,45) did not led to any changes in USP10 levels (Fig. S3E). Interestingly, reanalysis of a previously published proteomic dataset obtained from SH-SY5Y cells treated with TDP-43 fibrils that lead to endogenous TDP-43 aggregation (46), showed increased USP10 protein levels in cells with TDP-43 aggregates (Fig. S3F). Overall, these findings imply that, at least in specific contexts, TDP-43 can regulate USP10 levels.

### USP10 rescues TDP-43 dependent toxicity *in vivo*

As USP10 was upregulated in ALS and FTD patients, we finally tested the effects of USP10 upregulation on TDP-43 dependent toxicity in an *in vivo* animal model. To assess the effects of TDP-43 toxicity at the whole organism level, we produced a *Drosophila* model overexpressing human TDP-43 in glial cells, leading to major motor deficits associated with decreased survival. In parallel, we developed flies overexpressing human USP10 in glial cells, that did not show any overt motor or survival phenotypes and crossed them with the affected TDP-43 overexpressing strain. Remarkably, human USP10 co-expression with TDP-43 completely rescued the TDP-43 dependent motor deficits in the doubly transgenic flies (Fig. 3F) and potently reduced the associated mortality (Fig. 3G), without significantly affecting human TDP-43 transgene expression levels (Fig. S3G). Thus, USP10 can ameliorate TDP-43 induced toxicity *in vivo*.

## DISCUSSION

TDP-43 cytoplasmic aggregation coupled with its nuclear loss are the main pathological hallmarks defining TDP-43 proteinopathies, but how these aggregates are formed and what drives TDP-43 nuclear clearance have not yet been determined. Here, we propose USP10 as a new player in TDP-43 proteinopathies, regulating TDP-43 aggregation and functionality and reducing its toxicity.

Takahashi *et al*. recently described that USP10 promotes the clearance of TDP-43 cytoplasmic aggregates and the formation of TDP-43 aggresomes in cells treated with the proteasome inhibitor MG-132 (29). Some of their results are in contrast with our data, as we did not detect any induction of TDP-43 aggregation mediated by USP10 overexpression. The differences between both studies are likely explained by the different cellular models used, as in their study they mainly focussed on the role of USP10 on TDP-43 aggregation specifically under conditions of proteasome inhibition, which we do not assess here. Interestingly, they found a direct interaction between TDP-43 and USP10, which we have replicated (data not shown), that is mediated by TDP-43 RNA binding capacity and is likely to play a role in the wider functional connections between TDP-43 and USP10.

Although it is not yet clear how TDP-43 cytoplasmic foci might be formed and their possible relation with SG formation and resolution, we show here that USP10 can indeed modulate TDP-43 cytoplasmic aggregation in all models tested. In all the different paradigms used, USP10 inhibited TDP-43 aggregation irrespective of the cause of aggregation or its possible association with different co-aggregated components, such as whether it colocalizes with SG. This finding is compatible with previous reports proposing that TDP-43 colocalization with SG can be context dependent (47) and that TDP-43 aggregates could occur independent of SG (12). The levels and stoichiometry of G3BP1 and USP10 have been shown to be critical in modulating SG formation (16, 17, 22), with previous data showing that TDP-43 can control *G3BP1* levels through direct mRNA binding (18). Here, we have added another layer of complexity to the role of TDP-43 in modulating SG formation by showing that TDP-43 can, at least in some contexts, regulate USP10 levels. We note that there are multiple published TDP-43 mRNA binding sites in different *USP10* intronic regions (data not shown) that might play a role in this regulation. These findings places TDP-43, USP10 and G3BP1 in a functional hub that together can modulate SG formation, with TDP-43 regulating USP10 and G3BP1 levels, and USP10 in turn controlling TDP-43 levels, functionality and aggregation through a mechanism that requires USP10 interaction with G3BP1.

Moreover, we report that USP10, apart from its wider functions can also modulate transcriptome-wide alternative splicing. Interestingly, we identified a clear overlap between some USP10 modulated exons and a subset of TDP-43 splicing targets, supporting the newly identified functional connection. Further, we show that USP10 can ameliorate the splicing defects due to TDP-43 loss of function without affecting its expression levels (Fig. 2F and S2E). This, coupled with the inhibiting role of USP10 on TDP-43 aggregation, suggest that USP10 upregulation might be beneficial in TDP-43 proteinopathies by counteracting different aspects of TDP-43 toxicity. To directly test this hypothesis *in vivo*, we generated a *Drosophila* model leading to clear systemic toxicity and motor abnormalities upon TDP-43 overexpression, showing that USP10 upregulation can indeed potently rescue TDP-43 associated phenotypes at the whole organism level (Fig. 3F, G), identifying USP10 as a potential new therapeutic target in TDP-43 proteinopathies.

In patient material, we observed significantly increased USP10 levels in ALS and FTLD patients versus controls (Fig. 3C, D). These results are in contrast with the decrease in USP10 levels that we report in cellular models of acute TDP-43 downregulation (Fig. 3E), or the unaltered USP10 levels in chronic downregulation of *TARDBP* in i3 differentiated motor neurons (Fig. S3E). However, USP10 levels can increase in response to TDP-43 aggregation (Fig. S3F), suggesting that USP10 levels might increase in patients due to the presence of TDP-43 pathology. As our data from different cellular models, together with *Drosophila*, shows that USP10 overexpression could be protective against TDP-43 dependent toxicity, the upregulation seen in some patients might represent a cellular attempt to counteract the toxicity associated with TDP-43 proteinopathy. Moreover, the increase in USP10 levels might inhibit stress granule formation and modulate alternative splicing. Interestingly, recently, USP10 and G3BP1 protein levels have been found to be inversely correlated with TDP-43 pathological burden in postmortem human motor neurons (48), further supporting their functional connection. A number of studies have documented elevated levels of USP10 in other conditions, including glioblastoma and other cancers (49), as well as in Parkinson’s disease (PD) affected neurons (50), suggesting that USP10’s novel role as a splicing regulator may play a role in the pathophysiology of these disorders. Overall, from all the data presented, further investigations will be required to clarify the complex relation between TDP-43 and USP10 uncovered here, particularly to clarify the causes and consequences of USP10 upregulation in TDP-43 proteinopathies.

In conclusion, we describe a new functional interaction between USP10 and TDP-43, providing insights into some of the potential molecular mechanisms involved, with both proteins affecting different aspects of the biology of the other, including USP10 modulating TDP-43 aggregation and splicing function, both key molecular mechanisms in TDP-43 proteinopathies. Furthermore, we ascribe a new function to USP10 as a regulator of alternative splicing, with a clear overlap with some TDP-43 splicing targets, further supporting their functional connection. Crucially, we show that USP10 upregulation can rescue TDP-43 dependent toxicity in flies *in vivo*. All these data, together with the finding that USP10 can be altered in ALS and FTD patient material, suggest a possible involvement of USP10 in the pathogenesis of TDP-43 proteinopathies, giving potential clinical relevance to this newly identified functional interaction.

## MATERIALS AND METHODS

### Cell lines and cultures

All cell lines were cultured at 37° C with 5% CO_2_. HeLa, U2OS, Neuro2a were grown in DMEM media (31966, Gibco), whereas SH-SY5Y and SK-N-BE were grown in DMEM/F-12 (50:50), all supplemented with 10% FBS (10082147, Invitrogen) and 1% penicillin/streptomycin (10378016, Invitrogen).

The clover-TDP43^ΔNLS^ cell line was a kind gift from Don Cleveland (12). The inducible SK-N-BE cell line carrying a doxycycline inducible shRNA against TDP-43 has been published (51). For the induction of *TARDBP* gene silencing, doxycycline (1 µg/ml) (D9891, Sigma) was administered in the culture medium for 3-6 days, depending on the experiment. The inducible Hela cell line carrying the doxycycline inducible shRNA against TDP-43 was generated from a HeLa ATCC clone (CCL-2). Briefly, cells were transduced with an inducible system Tet-pLKO-puro (21915, Addgene) encoding an shRNA targeting *TARDBP* sequence (TRCN0000016038, SIGMA). Lentiviral particles were produced in the HEK293T cell line using psPAX2 (12260, Addgene) and pMD2.G (12259, Addgene) plasmids (both a kind gift from Dr Didier Trono). HeLa cells were transduced and selected with puromycin (1 μg/ml) for two weeks. A single clone was isolated and tested for KD efficiency (>80% reduction in protein expression) by doxycycline (200 ng/μl) exposure for three days.

iPSC i3LMN: The WTC11 iPSC line derived from a healthy human subject (GM25256) with stable integration of doxycycline-inducible expression of transcription factors NGN2, ISL1, and LHX3 (hNIL) at the CLYBL locus and stable integration of dCas9BFP-KRAB at the CLYBL locus was a gift from Michael Ward (45) and were maintained in E8 Flex Medium (Thermo) in Geltrex (Thermo) coated plates and passed with Versene solution (Thermo) or Accutase (Thermo) when 80% confluent. For induction of iPSCs to i3LMN, iPSCs were passaged with Accutase (Thermo) and plated into Geltrex-coated plates with induction media: DMEM/F12 with Glutamax (Thermo), 1x Non-essential amino acids (Thermo), 2 μg/ml doxycycline hyclate (Sigma), 1× N2 supplement (Thermo), and 0,098 μg/ml Compound E (Calbiochem). Media was changed daily for three days. On the second day cells were replated to poly-D-lysine and laminin coated plates. Differentiated neurons were maintained in motor neuron culture media: Neurobasal medium (Gibco) with 1x Non-essential amino acids (Thermo), 1x Glutamax (Thermo), 1x N2 supplement (Thermo), 1x N21 supplement (Thermo), 20 ng/mL BDNF (Peprotech), 20 ng/ml GDNF (Peprotech) 2 μg/ml doxycycline hyclate (Sigma), and 1 μg/mL laminin (Thermo) and half media changes were performed twice per week. For the induction of *TARDBP* gene silencing, we use the CRISPRi system (44). The iPSC line was transduced with a lentiviral construct expressing an sgRNA targeting *TARDBP* gene for 24h, with a subsequent selection by puromycin. Knockdown of *TARDBP* mRNA was robust in iPSCs and in i3LMN for several weeks after differentiation.

*Transfections* were performed with either jetPRIME (101000046, Polyplus), Lipofectamine 3000 (L3000015, Thermo Fisher) or electroporation with an Amaxa Nucleofector 2b device (AAB-1001, Lonza), all following manufacturer’s instructions. For all transfection experiments we used the same amount of DNA for each construct, keeping the total amount of transfected DNA constant with EV. Cells were transfected and harvested after 24 hours, unless otherwise stated.

#### Cellular stress treatments

To induce TDP-43 cytoplasmic localization and SG in SH-SY5Y, coverslip-plated cells were treated with 500 µM NaAsO_2_ for 40 minutes, followed by incubation with 600 mM sorbitol for 40 minutes. Cells were then left to recover with fresh media for 1 hour and coverslips processed for microscopy. To induce SG in U2OS cells, 250 µM NaAsO_2_ was used for 80 minutes. To induce TDP-43 cytoplasmic aggregates in the clover-TDP43^ΔNLS^ cell line, TDP-43 cytoplasmic localization was induced by doxycycline treatment (1 µg/ml doxycyclin for 24 hours), followed by 250 µM NaAsO_2_ for 80 minutes.

*Small interfering RNAs (siRNAs)* specific to human TDP-43 (siTDP43: GCAAAGCCAAGAUGAGCCUTTdTdT), human USP10 (siUSP10: CCCUGAUGGUAUCACUAAAGAdTdT) or negative control luciferase (siLuc: CGUACGCGGAAUACUUCGAUU dTdT), were transfected into U2OS or HeLa cells using Lipofectamine RNAiMAX reagents (Invitrogen) according to the manufacturer’s protocol. When a combination of downregulation and overexpression was performed a first round of downregulation was performed on day, followed by a second round on day 2. Then, on day 3, cells were transfected for overexpression using lipofectamine 3000, and harvested on the next day.

### Constructs

The GFP-tagged TDP-43-WT and TDP-43^ΔNLS^ constructs were a kind gift from the laboratory of Leonard Petrucelli (52). The CFTR C155T TG11T5 minigene was a kind gift from Emanuele Buratti (53,54). The USP10 WT and USP10^C424A^ construct were kindly provided by Lorenza Penengo (University of Zurich, Switzerland). The USP10^F10A^ mutant was made using site directed mutagenesis following manufacturer’s instructions (Agilent). All constructs were amplified in commercial *E. coli* lines and plasmids purified (740410.50, Macherey Nagel) according to the manufacturer’s instructions.

### Antibodies

The following antibodies and concentrations were used for immunoblot and immunofluorescence staining: Rabbit anti TDP-43 (WB 1:1000, IF 1:400, Protein Tech, 10782-2-AP); mouse anti TDP-43 (WB 1:5000, abcam, ab104223); mouse anti TDP-43 (IF 1:200, R&D systems, MAB37778); mouse anti pTDP-43 (IF 1:3000, Cosmobio, CAC-TIP-PTD-M01); mouse anti G3BP1 (IF 1:300, Protein Tech, 66486-1-Ig); rabbit anti G3BP1 (WB: 1:2000, IF 1:300, Protein Tech, 13057-2-AP); rabbit anti G3BP2 (IF 1:300, Protein Tech, 16276-1-AP); rabbit anti USP10 (WB 1:1000, IF 1:500, Sigma-Aldrich, HPA006731); rabbit anti USP10 (WB 1:1000, made in house from the Freire laboratory); rabbit anti USP10 (IF 1:200, Bethyl, A300-901A); mouse anti Tubulin (WB 1:10000, Protein Tech, 11224-1-AP); mouse anti β-Tubulin (WB 1:10000, Protein Tech, 66240-1-Ig); mouse anti GAPDH (WB 1:250, Santa Cruz Biotechnology, sc-32233); anti mouse Alexa Fluor 488 (IF secondary 1:600, Thermo Fisher, Z25002); anti rabbit Alexa Fluor 488 (IF secondary 1:600, Thermo Fisher, Z25302); anti mouse Alexa Fluor 568 (IF secondary 1:600, Thermo Fisher, Z25006); anti rabbit Alexa Fluor 568 (IF secondary 1:600, Thermo Fisher, Z25306). The antibody against human USP10 was generated by injecting rabbits with a His-tagged antigen (amino acids 1-350). The antigen was obtained by cloning the corresponding cDNA in pET-30 (Novagen), followed by expression in E. coli and purification with a Ni-NTA resin (Qiagen) following manufacturers recommendations.

### Patient material

We obtained frontal cortex brain tissue from the UCL Queen Square Brain Bank, separated FTLD cases according to the subtype of TDP-43 pathology and a control group of deceased patients with no pathologies described. Lysates from ALS motor cortex and controls came from the Bellvitge Hospital (Barcelona). All biological samples comply with the legal requirements of consent, collection, use and storage.

### Immunofluorescence

Cells were grown in plates containing sterile glass coverslips, fixed with 4 % PFA (P6148, Sigma-Aldrich), permeabilised using 0.5% Triton detergent (85111, Thermo Fisher) and blocked with BSA (422381B, VWR). The samples were then incubated overnight with primary antibodies in BSA, and then with appropriate secondary antibodies. Slides were mounted using ProLong™ (P36962, Thermo Fisher) and samples visualised on a Zeiss Axio Observer fluorescence microscope, with images analysed using Image J software 50.

The immunohistochemistry experiments from patient material were carried out on 20 μm thick paraffin-embedded sections obtained from the Queen Square Brain Bank for Neurological Disorders (University College of London). Antigenic unmasking was performed using the Dako PT Link PT100 Slide Stainer (Swerdlick Medical Systems), prior to permeabilization with 0.5% Triton. Immunofluorescence or DAB was used for visualized (15454629, Thermo Fisher) using a Leica DMi1 light microscope.

For the detection and quantification of TDP-43-positive aggregates and SG, pictures of 50-100 cells from random fields were taken with a Zeiss Axio Observer fluorescence microscope, and the images taken analysed using Image J software. Aggregates were analysed with the Aggrecount (55) Image J plugin. The data collected were the number of aggregates per cell, the number of cells with aggregates and the size of the aggregates. To consider that a cell had TDP-43 inclusions or SG, cells with more than two TDP-43 inclusions/SG larger than 0.5-1 µm^2^ were taken as positive. For colocalization studies, a line profile analysis was performed and the quantification was done using JaCoP in Image J giving the Pearson coefficient and the mass centre coincidence (56).

### Protein extractions

From cell lines, proteins were extracted by resuspending cells from 6-well plates in 100 μl of RIPA or 7M Urea Buffer containing protease inhibitor (11873580001, Roche) and, if necessary, phosphatase inhibitor (04906837001, Roche). Cell lysates were centrifuged at >14 000 x g for 15 minutes, discarding the sediment. Supernatant was then sonicated using an ultrasonic homogeniser (UP50H, Hielscher) for 15 seconds. Protein concentration in the soluble extract was quantified by using the BCA method (Thermo Scientific). For post-mortem material from FTD patients, 13mg of frozen sample was weighed, to which 0.26 ml of 7M Urea buffer was added and homogenised using a rotary homogeniser (BK-HG, Biobase). The samples were then sonicated using an ultrasonic homogeniser (UP50H, Hielscher) for 15 seconds. Samples were frozen at -80°C until use.

### Immunoblots

Cell lysates were resuspended in Laemmli buffer and heated at 95°C for 5 minutes. Then, proteins were separated by 10% SDS polyacrylamide gel electrophoresis (SDS-PAGE) using a Mini-PROTEAN kit (1658001FC, Bio-Rad). Between 20 and 50 μg of protein per well was loaded. After the separation, proteins were transferred onto a nitrocellulose (1620115, BioRad) or PVDF (1620177, Bio-Rad) membrane, that was blocked with either BSA or milk powder. The primary antibodies and concentrations used are listed in the antibodies section.

Bands were detected via ECL (1705060, Bio-Rad), viewed on a ImageQuant™ LAS 4000 chemiluminescence reader (GE Healthcare), and analysed by densitometry using the Image Quant TL software.

### RNA isolation, RT–PCR and qPCR

RNA extractions were carried out using TRIzol (15596-026, Invitrogen), according to manufacturer’s instructions. For complementary DNA synthesis, the qScript cDNA synthesis kit (95047, QuantaBio) was used, using as a template 100-500 ng of RNA. RT-PCR for splicing events was conducted using 2X PCR Master Mix (11816843, Thermo Fisher) with specific primers (see below) spanning the differentially expressed exon. Products were electrophoresed on 2.5% agarose gels containing ethidium bromide and visualized on an automated image detection system (Gel Doc EZ, BioRad). The BioRad Image Lab software was used to quantify the bands. Results were analysed using the percentage of exclusion (PSE), calculated dividing the intensity of the shorter exon exclusion band by the sum of both exon inclusion and exon exclusion bands. Primers used here were:

*RABGEF exon 16*: CTGCAGACAAAGCACACACGAT and GCACAGAGGAGATCCGATTGAT

*RWDD1 exon 2*: GGGCCACGATGACAGATTAC and AGGAGGGTTGGATCAATGTG

*CFTR minigene*: CAACTTCAAGCTCCTAAGCCACTGC and TAGGATCCGGTCACCAGGAAGTTGGTTAAATCA

qPCR was used to quantify mRNA expression using specific primers (see below). SYBR Green (PB20.11-05, PCR Biosystems) was used on a real-time thermal cycler (LightCycler® 480, Roche). *S16* or *HPRT* were used as internal control gene. The levels of messenger RNA were assessed by calculating ΔΔCT which shows the relative expression change (fold change), normalised to the housekeeping gene. For all samples analysed, 3 technical replicates and 3 biological replicates were performed. The primers:

*GFP*: AAGCAGAAGAACGGCATCAAG and TCAGGTAGTGGTTGTCGGGCA

*USP10*: GGGTGCCAGAAGCTTATCAA and CACTGTTGCCGTGATGGTAG

*TARDBP (exon 1-2)*: CTGCTTCGGTGTCCCTGT and ATGGGCTCATCGTTCTCATC

*TARDBP (exon 5-6)*: ATGACTGAGGATGAGCTGCG and CACAAAGAGACTGCGCAATCT

*S16*: AAGTCTTCGGACGCAAGAAA and TGCCCAGAAGCAGAACAG

*HPRT*: TTTGCTGACCTGCTGCTGGATTAC and TTGACCATCTTTGGATTATACTGC

### RNA sequencing analysis

RNA was quality checked using a Tapestation (Agilent) prior to submission to the UCL Genomics facility; sequencing libraries were prepared with polyA enrichment and sequenced with 150 nt paired reads.

Raw sequences (in fastq format) were trimmed using Fastp with the parameter “qualified_quality_phred: 10”, and aligned using STAR (v2.7.0f) to the Grch38 with gene models from GENCODE v40 (57–59).

The STAR outputs are BAM files, containing the counts and coordinates for all splice junctions found in the sample, and alignment metadata including the genomic location where a read is mapped to, read sequence, and quality score. Then, using samtools, BAM files were sorted and indexed by the read coordinates location (60).

Trimmed fastqc files were aligned to the transcriptome using Salmon (v1.5.1), outputting isoform-specific counts used for differential gene expression, performed using DeSEQ2 without covariates, using an index built from GENCODE v40 (61,62). The DESeq2 median of ratios, which controls for both sequencing depth and RNA composition, was used to normalise gene counts. Differential expression was defined at a Benjamini–Hochberg false discovery rate < 0.1. We kept genes with at least 5 counts per million in more than 2 samples.

Splicing analysis was performed using Majiq (v2.1) on STAR-aligned and sorted BAMs and the Grch38 reference genomes. Cryptic splicing was defined as junctions with Ψ (PSI, percent spliced in) < 5% in the control samples and ΔΨ > 10% between groups, while skiptic splicing as junctions with Ψ > 95% in the control samples and ΔΨ < - 10% between groups. Other junctions were classified as differentially spliced when the absolute value of ΔΨ was above 10% between groups.

The alignment and splicing pipelines are implemented in Snakemake (v5.5.4), a workflow management software that allows reliable, reproducible, and scalable analysis, and run on the UCL high-performance cluster (HPC) (63). DESeq2 and data visualisation were run using R with the packages tidyverse, rstatix, and eulerr, and custom scripts available upon request.

### USP10 analysis from published datasets

The mass spectrometry proteomics data from Scialò C et al (46), was downloaded from the ProteomeXchange Consortium repository (64) under the dataset identifier PXD058981. From the report.tsv file generated by DIA-NN, which contains precursor ion abundances for each raw file, protein abundances were obtained by first aggregating precursor abundances to peptidoform abundances, followed by estimation via Tukey’s median polish using the R package prolfqua (65). Protein abundances were transformed using robscale prior to fitting linear models, and expressed relative to negative controls, which were set to a value of 1. Data are shown as mean with s.d. as indicated in the figure legends. Statistical significance was assessed by t-test.

The RNA-sequencing data from Scialò C et al (46), have been downloaded from GEO database (66) under accession number GSE285224. The normalized counts were expressed relative to negative controls, which were set to a value of 1. Data are shown as mean with s.d. as indicated in the figure legends. Statistical significance was assessed by t-test.

### Drosophila models

Different *Drosophila* strains were obtained from Vienna Stock Center (VDRC, Austria) and Bloomington Stock Centre (BDSC, Indiana-USA). REPO was used as the driver for conditional expression in glial cells (BDSC, #7415), that was crossed with strains carrying sequences to express human TDP-43 (BDSC, #79587) and USP10, the later one was newly generated using Bestgene Inc. via their embryo injection service, with the strain selected expressing the highest amount of USP10 measured by RT-PCR.

The analysed experimental genotypes for glial expression were (G-E)-(UAS-hTDP43///REPO-Gal4) and (G-E)-(UAS-hUSP10///REPO-Gal4). Stocking flies were housed at 22 °C, 70% humidity and 12 h/12 h light/darkness cycle and experimental flies at 26 °C, 70% humidity and 12 h/12 h light/darkness cycle.

#### Drosophila functional assays

For longevity tests, 50 adult female flies (five per tube) were selected from each strain. Dead flies were counted every 2 days and tubes were replaced every 7 days. Mantel-Cox method was used to analyse the results. For locomotor activity, adult flies were tested on day 7 in groups of five (the same groups as in the longevity test). They were placed in a tube with a line drawn outside the tube at a height of 8 cm from the bottom. The number of flies crossing the line in 10s was counted (3 trials per tube). The results were analysed using t-student analysis. Adult pharate survival was expressed as a percentage of adult flies counted over the total number of pupae in each tube, where adult pharate survival of control flies were approximately 100%.

#### Drosohila RNA extraction and quantitative real time PCR

Total RNA was extracted from 2-4 independent experimental pools using the RNeasy Mini Kit (ref. 74104, QIAGEN). Each pool consisted of 10 thoraces, 20 brains or 30 ventral nerve cords (VNCs), depending on the tissue to be analysed. Reverse transcription was performed using the SuperScript VILO cDNA Synthesis Kit (Thermo Fisher) according to the manufacturer instructions.

Quantitative real-time PCR was performed on a CFX384 Touch Real-Time PCR Detection System (Bio-Rad) using Powe SYBR Green Master Mix (Thermo Fisher), 300 nM of primer pair, and 10 ng of cDNA. Tubulin (Tub) was used as a housekeeping gene. The 2-ΔΔCt method was used for relative quantification. The primers sequences used for genotype confirmation were pre-designed and obtained from Sigma-Aldrich: Tbph - DMEL_TBPH_8, TARDBP – H_TARDBP_1.

### Statistics

Results were analysed using GraphPad Prism 8.0.2 software. Data are generally presented as the mean accompanied by the standard error (Mean±SEM). Test and sample sizes used are generally specified in each figure. For parametric samples, t-test and/or ANOVA was performed. For non-parametric samples, the Mann-Whitney U test or Kluskal Wallis were used. Statistical significance is attributed at a p value <0.05.

## Acknowledgements

We thank Professor Elizabeth MC Fisher for critical reading of the manuscript. We are grateful to Professors Don Cleveland, Leonard Petrucelli, Sandrine Da Cruz, Michael Ward and Lorenza Penengo for sharing critical reagents.

