## Supplementary figures and legends for "A functional interaction between TDP-43 and USP10 reveals USP10 dysfunction in TDP-43 proteinopathies"

### Figure S1

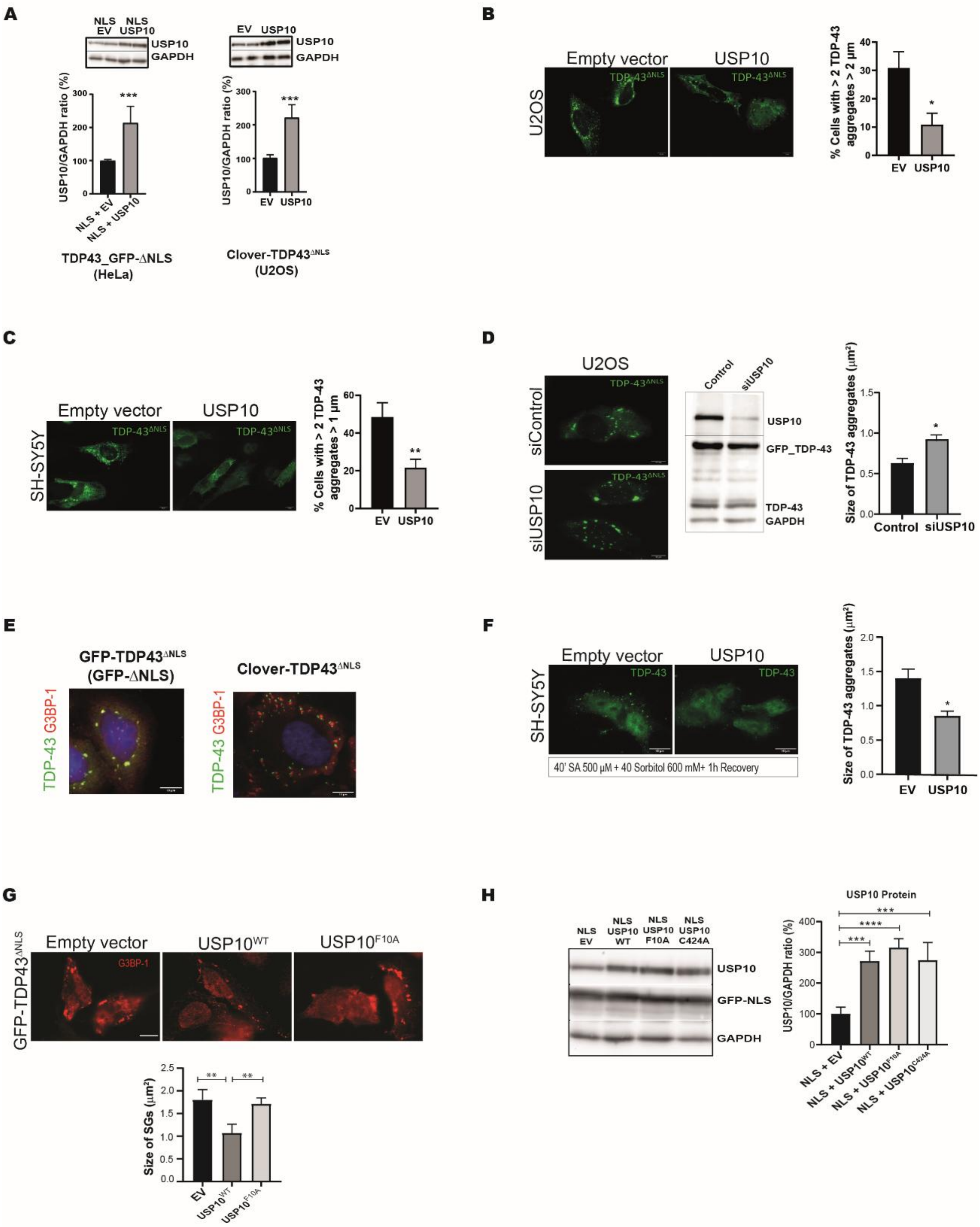

A
NLS EV
NLS USP10
USP10
GAPDH
USP10/GAPDH ratio (%)
***
NLS + EV
NLS + USP10
TDP43_GFP-ΔNLS (HeLa)
EV
USP10
Clover-TDP43ΔNLS (U2OS)
B
Empty vector
USP10
U2OS
TDP-43ΔNLS
% Cells with > 2 TDP-43 aggregates > 2 µm
C
SH-SY5Y
% Cells with > 2 TDP-43 aggregates > 1 µm
D
siControl
siUSP10
Control
GFP_TDP-43
TDP-43
Size of TDP-43 aggregates (µm²)
E
GFP-TDP43ΔNLS (GFP-ΔNLS)
Clover-TDP43ΔNLS
TDP-43 G3BP-1
F
40' SA 500 µM + 40 Sorbitol 600 mM+ 1h Recovery
G
GFP-TDP43ΔNLS
USP10WT
USP10F10A
G3BP-1
Size of SGs (µm²)
H
USP10 Protein
NLS USP10 WT
NLS USP10 F10A
NLS USP10 C424A
GFP-NLS
NLS + USP10WT
NLS + USP10F10A
NLS + USP10C424A
****

**Supplementary Figure 1 (S1). (A)** Representative image of immunoblot in HeLa cells transfected with the indicated constructions. USP10$^{WT}$, USP10$^{F10A}$ and USP10$^{C424A}$ are expressed at the same levels in conditions of TDP-43$^{\Delta NLS}$ co-transfection. **(B)** Representative fluorescence images showing TDP-43 aggregates in U2OS and **(C)** SH-SY5Y cells, 24 hours after co-transfection with GFP_TDP-43$^{\Delta NLS}$ and USP10 or an EV. Scale bar: 10 µm. Percentage of U2OS and SH-SY5Y cells with more than two TDP-43 aggregates bigger than 1 µm after co-transfection of TDP-43$^{\Delta NLS}$ with USP10 compared to EV control by Mann-Whitney U test. The quantification shown corresponds to one independent experiment from a total of two. At least 50 cells per group were analysed. *p<0,01; **p<0,001. Error bars show the SEM for all figure panels. **(D)** Representative images showing TDP-43 aggregates in U2OS cells 24 hours after co-transfection of GFP_TDP-43$^{\Delta NLS}$ after 72 hours of USP10 downregulation with siRNA. Scale bar: 10 µm. The quantification of the area and percentage of U2OS cells with more than two TDP-43 aggregates bigger than 1 µm after USP10 downregulation with siRNA during 72 h. At least 50 cells per group from 2 independent experiments were analysed; *p<0.05 analysed by Mann-Whitney U test compared to siLuciferase control. **(E)** Colocalization of TDP-43 with G3BP1 in HeLa cells transfected with GFP_TDP-43$^{\Delta NLS}$, Pearson´s correlation coefficient of the colocalization between TDP-43 and G3BP1 (Pearson=0.93), and U2OS TDP-43$^{\Delta NLS}$-Clover after induction with doxycycline and stress treatment (Pearson=0.09), The quantification shown corresponds to one experiment. Representative images of fluorescence microscopy of the two cellular models, showing TDP-43 aggregates (green) and SG (labelled with anti-G3BP1 primary antibody in red). Scale bar: 10 µm. **(F)** Representative fluorescence images showing TDP-43 aggregates (labelled with anti-TDP-43 primary antibody in green) in SH-SY5Y cells 24 hours after transfection with an EV or USP10. TDP-43 aggregation was induced by incubating the cells consecutively with 500 µM NaAsO2 (40 minutes), 600 mM sorbitol (40 minutes) with 1 hour recovery after the treatment with fresh media. Scale bar: 10 µm. Quantification of the area and percentage of SH-SY5Y cells with more than two TDP-43 aggregates bigger than 0.6 µm after transfection with USP10 when compared to the EV. *p<0.05, analysed by Mann-Whitney U test. At least 50 cells per group from 2 independent experiments were analysed. **(G)** Representative fluorescence image and quantification of SG size in HeLa cells after co-transfection with GFP_TDP-43$^{\Delta NLS}$ and either USP10, USP10$^{F10A}$ or an empty vector (EV) control. The quantification shown corresponds to one independent experiment from a total of two. At least 50 cells were analysed per experiment. NS: Not significant; **p<0.01, ***p<0.001 analysed by Mann-Whitney U test compared to EV. **(H)** Representative image of immunoblot in U2OS TDP-43$^{\Delta NLS}$-Clover cells transfected with either an empty vector or USP10$^{WT}$, USP10$^{F10A}$ and USP10$^{C424A}$, showing that the three constructs are similarly overexpressed after transfection in this cellular model.

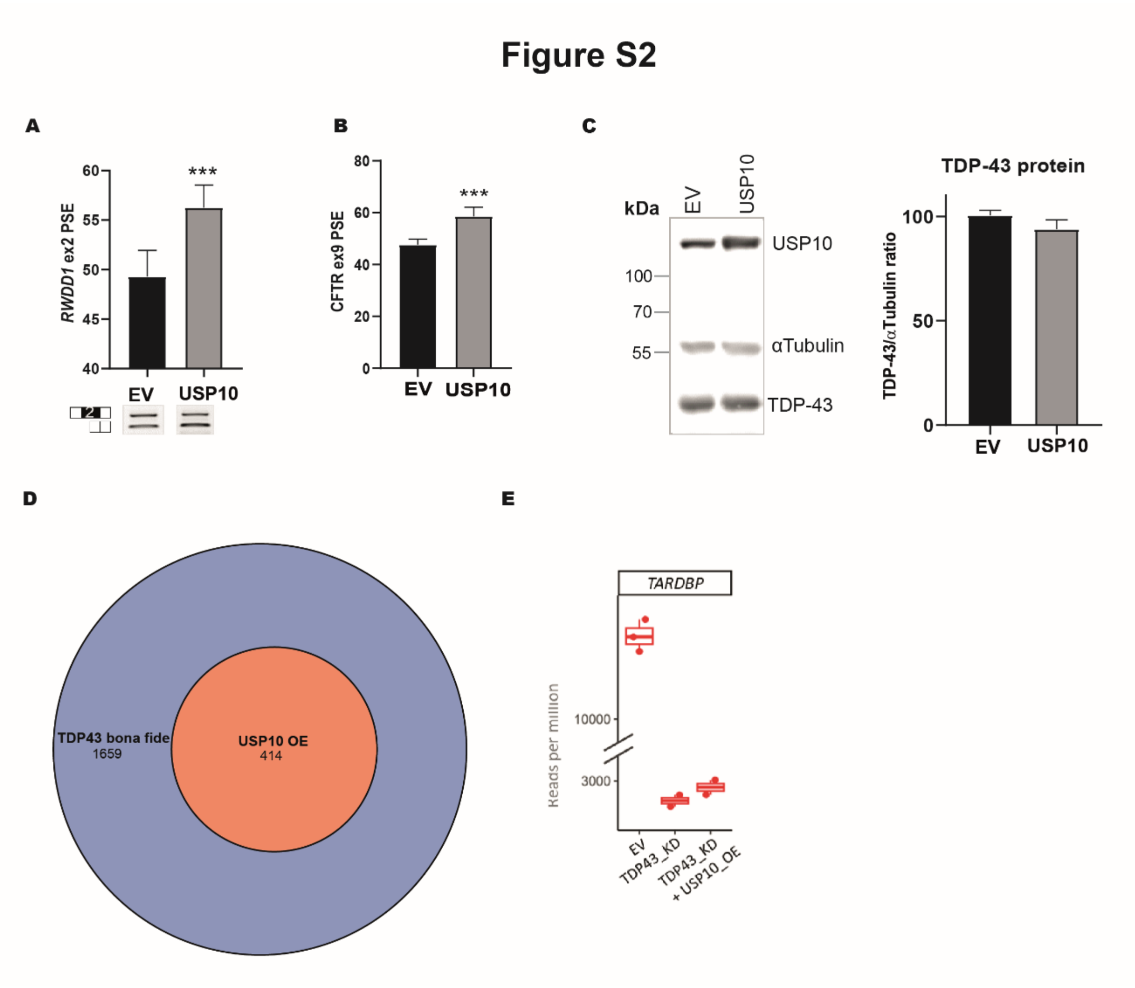


**Supplementary Figure 2 (S2). (A)** Representative image and quantification of the PSE of *RWDD1* exon 2. HeLa cells were transfected with either an EV or USP10. Following RT-PCR, the PSE was quantified and compared via one-way ANOVA. **(B)** Representative image and quantification of the PSE of *CFTR* exon 9. U2OS cells were co-transfected with the *CFTR* minigene and either an EV or USP10. Following RT-PCR, the PSE was quantified, and all groups compared to the EV via Mann Whitney test (*$p<0.05$) and error bars represent SEM. N=3, from 1 experiment. (**C**) Representative image of immunoblot in U2OS cells transfected with USP10 or EV. Specific primary antibodies against USP10, Tubulin, and TDP-43 were used. TDP-43 levels were quantified via densitometry normalized to their loading control (Tubulin). No changes in TDP-43 protein levels were detected following USP10 overexpression compared to the EV by Mann-Whitney U-test ($p=0.8$). N=9, from 3 independent experiments. Error bars show the SEM for all panels. (**D**) Venn diagram representing the overlap between *bona fide* TDP-43 splicing targets moving in the direction of TDP-43 overexpression upon USP10 overexpression **(E)** Quantification of *TARDBP* transcript by RNA-seq analysis in reads per million, after either TDP-43 downregulation or TDP-43 downregulation + USP10 overexpression in HeLa cells, compared to its expression when transfecting HeLa cells with an Empty Vector.

### Figure S3

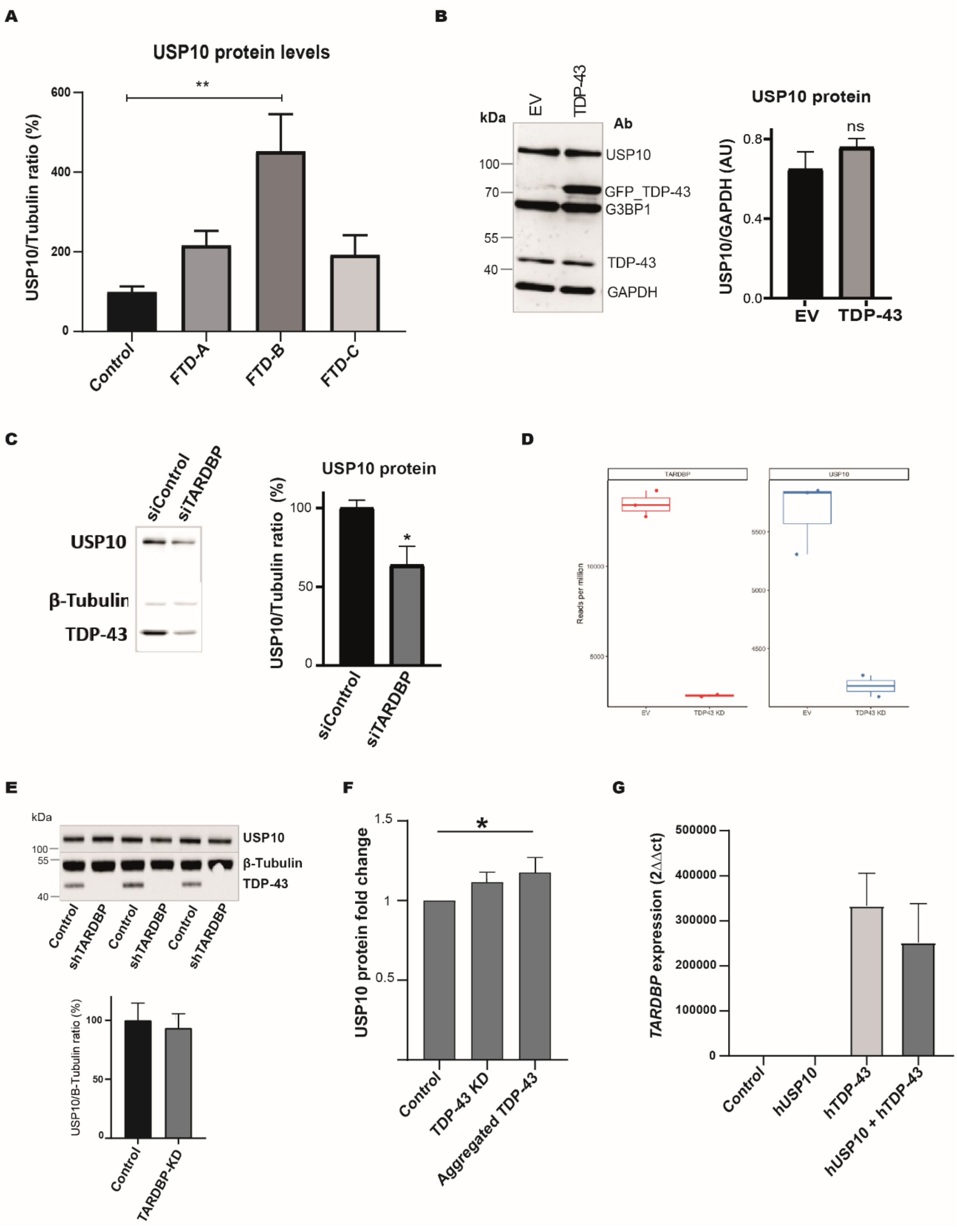

A
USP10 protein levels
USP10/Tubulin ratio (%)
**
Control
FTD-A
FTD-B
FTD-C
B
kDa
EV
TDP-43
Ab
USP10
GFP_TDP-43
G3BP1
TDP-43
GAPDH
USP10 protein
USP10/GAPDH (AU)
ns
C
siControl
siTARDBP
USP10
β-Tubulin
TDP-43
USP10 protein
USP10/Tubulin ratio (%)
*
D
TARDBP
USP10
Reads per million
EV
TDP43 KD
E
kDa
USP10
β-Tubulin
TDP-43
Control
shTARDBP
USP10/Β-Tubulin ratio (%)
Control
TARDBP-KD
F
USP10 protein fold change
*
Control
TDP-43 KD
Aggregated TDP-43
G
TARDBP expression (2ΔΔct)
Control
hUSP10
hTDP-43
hUSP10 + hTDP-43

**Supplementary Figure 3 (S3). (A)** Representative image of immunoblots from frontal cortex protein lysates of FTD patients and controls and quantification of USP10 bands by densitometry normalized to their loading control (Tubulin), separating the patient samples by the different FTLD-TDP pathology subtypes. USP10 levels were particularly increased in samples from patients with TDP-43 type B pathology ($p<0.01$). Samples from 3 patients per pathology subtype and 3 controls were used and compared pairwise via one-way ANOVA to the control group. *$p<0.05$, **$p<0.01$, ***$p<0.001$ for all panels. Error bars represent SEM for all figure (**B**) Representative image of immunoblot and quantification of HeLa cells transfected with TDP-43 or EV. Quantification of USP10 bands by densitometry was normalized to their loading control (GAPDH). No changes in USP10 protein levels were detected following TDP-43 overexpression ($p=0.486$ by Mann-Whitney U-test). N=8, from 3 independent experiments. The quantification shown corresponds to one experiment. (**C**) Representative image of immunoblot and quantification of HeLa cells downregulated during 72 h. with either siTDP-43 or a negative control (siControl). Quantification of USP10 bands by densitometry was normalized to their loading control (β-Tubulin), showing reduced USP10 protein levels upon TDP-43 downregulation when compared to the siControl by unpaired T test to control. N=3, from 1 independent experiment. (**D**) Quantification (reads per million) of *TARDBP* and *USP10* transcripts by RNA-seq analysis after TDP-43 downregulation in HeLa cells, compared to the EV. **(E)** Representative image of immunoblot and quantification of i3LMN iPSCs differentiated onto lower motor neurons, CRISPRi-mediated knockdown for 14 days of TDP-43 compared to controls. Quantification of USP10 bands by densitometry was normalized to their loading control (β-Tubulin). The quantification shown corresponds to 3 independent experiments, corroborated by another independent experiment by T test ($p= 0.58$). **(F)** Quantification of USP10 expression levels from proteomics data from Scialo *et al* of SH-SY5Y cells treated with exogenous TDP-43 fibrils. Aggregated TDP-43 bar corresponds to the ATTO group from the publication. Comparison between the control group and the TDP-43 aggregated by T test ($p= 0.03$). **(G)** TDP-43 overexpression transgene levels quantified via qRT-PCR from RNA obtained from the fly brains of the selected genotypes show no differences in TDP-43 transgene expression levels upon USP10 co-overexpression ($p=0.08$). Samples from 3 flies per group were used and compared pairwise via one-way ANOVA to the TDP-43 overexpression group.
